# Effective Rapid Blood Perfusion in *Xenopus*

**DOI:** 10.1101/2023.02.01.526649

**Authors:** Rachael A Jonas-Closs, Leonid Peshkin

## Abstract

*Xenopus* has been a powerful model organism for understanding vertebrate development and disease for over a hundred years. Here we define a rapid blood perfusion protocol in *Xenopus* aimed at a consistent and drastic reduction of blood across tissues. Perfusion is done by inserting a needle directly into the ventricle and pumping heparin in PBS through the vascular system. The whole procedure should take about 10 minutes per frog. Blood is dominated by a few highly abundant proteins and cell types which create numerous issues by masking most other molecules and cell types of interest. Reproducible characterization of adult *Xenopus* tissues with quantitative proteomics and single cell transcriptomics will gain from applying this protocol prior to organ dissections defined in companion papers. The procedure is aimed at standardization of practice across the animals of different gender, age and *Xenopus* species, specifically *X.laevis* and *X.tropicalis*.

**SUMMARY:** An effective rapid blood perfusion protocol to prepare tissue samples for transcriptomics and proteomics studies.

## Introduction

We set out to develop in *Xenopus* an effective blood perfusion protocol to parallel these developed in mice [10] and axolotl [4] while making the technique speed the top priority to ensure sample freshness. The protocol conservatively takes less than 10 mins per animal in X. *tropicalis* and less than 15 minutes per *X.laevis* animal. The secondary priorities were the ease of replication and the use of easily acquired equipment so that the technique could be utilized by the majority of *Xenopus* labs and high-quality samples can be shared widely.

*Xenopus* frogs are used widely in biomedical research to study fundamental biological and pathological processes conserved across species. The international community effectively uses *Xenopus* to gain a deeper understanding of human disease through in-depth disease modeling and molecular analysis of disease-related gene function. The numerous advantages of *Xenopus* as the animal model make them invaluable tools to study the molecular basis of human development and disease and include: large oocyte and embryo size, high fecundity, ease of housing, rapid external development, and ease of genomic manipulation. This tetrapod has a closer evolutionary relationship with mammals than other aquatic models for having lungs, a three-chambered heart, and limbs with digits. *Xenopus* has been estimated to share ~80% of the identified human disease genes [5].

Compared to popular mammalian models, *Xenopus* is a rapid, cost-effective model with the ease of morpholino knock-down, and availability of efficient transgenics and targeted gene mutations using CRISPR [6]. Such methods as quantitative mass spectrometry and single-cell proteomics have been successfully applied to Xenopus embryos [7,8] but a cell type atlas in adult *Xenopus* shows that the composition of most tissues is dominated by blood specific cell types [9]. By developing a technique that exsanguinates tissue at a rapid rate and uses chilled media, sample quality is improved without compromising the sample freshness. This is particularly important for applications, where we seek to profile mRNA and protein expression before any stress response or necrotic signatures, get pronounced.

### Protocol

All *Xenopus* experimentation discussed in this paper was performed in accordance with the rules and regulations of the Harvard Medical School IACUC (Institutional Animal Care and Use Committee). (IS 00001365_3). Though the primary method of euthanasia described is deemed an acceptable technique for euthanasia by the AVMA [11] it has not been found to lead to the cessation of heartbeat [12]. Even the frequently used secondary method of damaging the brain and spine through double pithing does not prevent the *Xenopus* heart from beating, nor does removing the heart from the animal. Exsanguination of anesthetized animals is a less usable but more effective method for successful euthanasia. It serves this protocol that the heart rate of animals continues through primary euthanasia and that perfusion is itself a secondary method, through exsanguination. This technique should be approved by institutions prior to being attempted.

### Preparation

1. Ensure that the described euthanasia techniques are approved by your institution. Prepare a solution of 5g/L MS-222 (tricaine methanesulfonate) and 5g/L sodium bicarbonate. Check the pH to ensure that it is ≥7.
2. Perform primary euthanasia by placing the *Xenopus* in this solution for 1 hour.
3. 15 minutes into euthanasia confirm that the *Xenopus* has lost its pain response by pinching the foot. Do not continue if the animal is reactive.
4. Using a 31 gauge needle inject 100uL of 180 units/mL heparin in PBS into the musculature of each forelimb.
5. Blunt your perfusion needle by trimming off the tip with wire cutters (Fig. 1) [2]. This will reduce the likelihood of the needle perforating through the ventricle if shifted.
6. Prepare the pump by attaching the trimmed needle and circulating 54 units/mL heparin in PBS. Ensure that all air bubbles have been purged from the line to eliminate the possibility of air emboli which lead to decreased perfusion efficiency or failure (troubleshoot 4). The perfusion media should be kept on ice for the duration of the procedure.
7. If the perfusion pump is programmable then calibrate it with the needle in place. If your pump is not programmable, with the needle in place, measure the pumped volume of media under the different settings to determine which settings are closest to 5mL/min and 10mL/min.
8. Place the dissection surface (tray or foam sheet) at an incline within a secondary container or otherwise arrange it to facilitate drainage.
9. Once primary euthanasia has been completed remove the frog and recheck the loss of pain response by performing a foot pinch. Place it on its dorsum and pin down each limb. (Fig. 2)
10. Using dissection scissors cut up the midline and then laterally, making 2 flaps. (Fig. 3)
11. Use forceps to grasp the linea alba and pull it away from the coelomic cavity (also referred to as the pleuroperitoneal cavity). Carefully use scissors to cut the peritoneum. Make 2 additional flaps and cut or pin all flaps out of the way.
12. Use dissection scissors to cut up through the coracoid bones and cut away excess tissue to gain better access to the heart.
13. It is likely that the heart will still be beating. Perfusion will function as the secondary method of euthanasia. If the heart has stopped beating prior to perfusion this should be noted with your samples.

**Figure 1.**
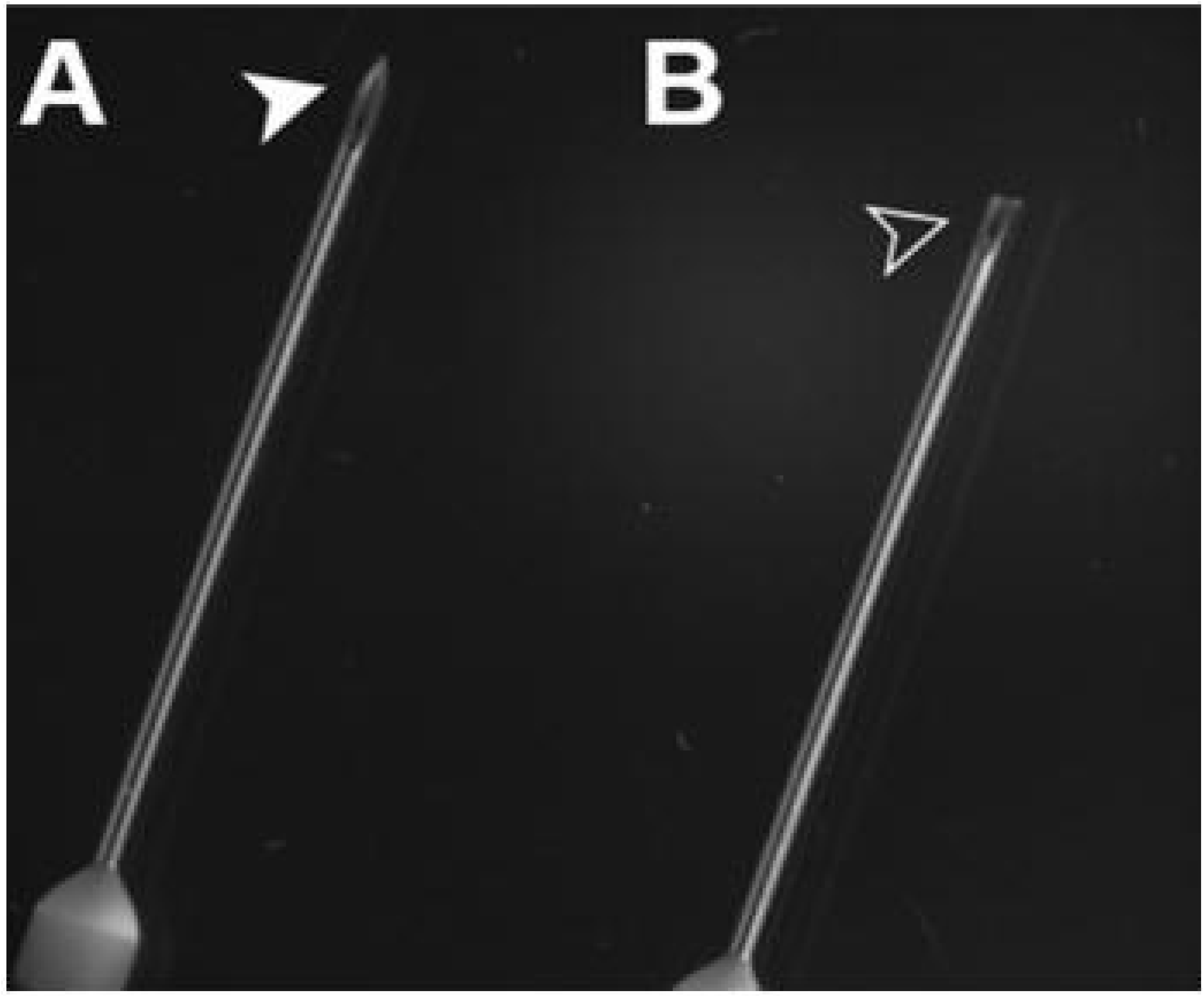
Untrimmed and trimmed needles [2]. Using wire clippers, blunt your needle by cutting off its tip. It will be sharp enough to pierce the heart, yet perforating the ventricle will be less likely in the event of human error.

**Figure 2.**
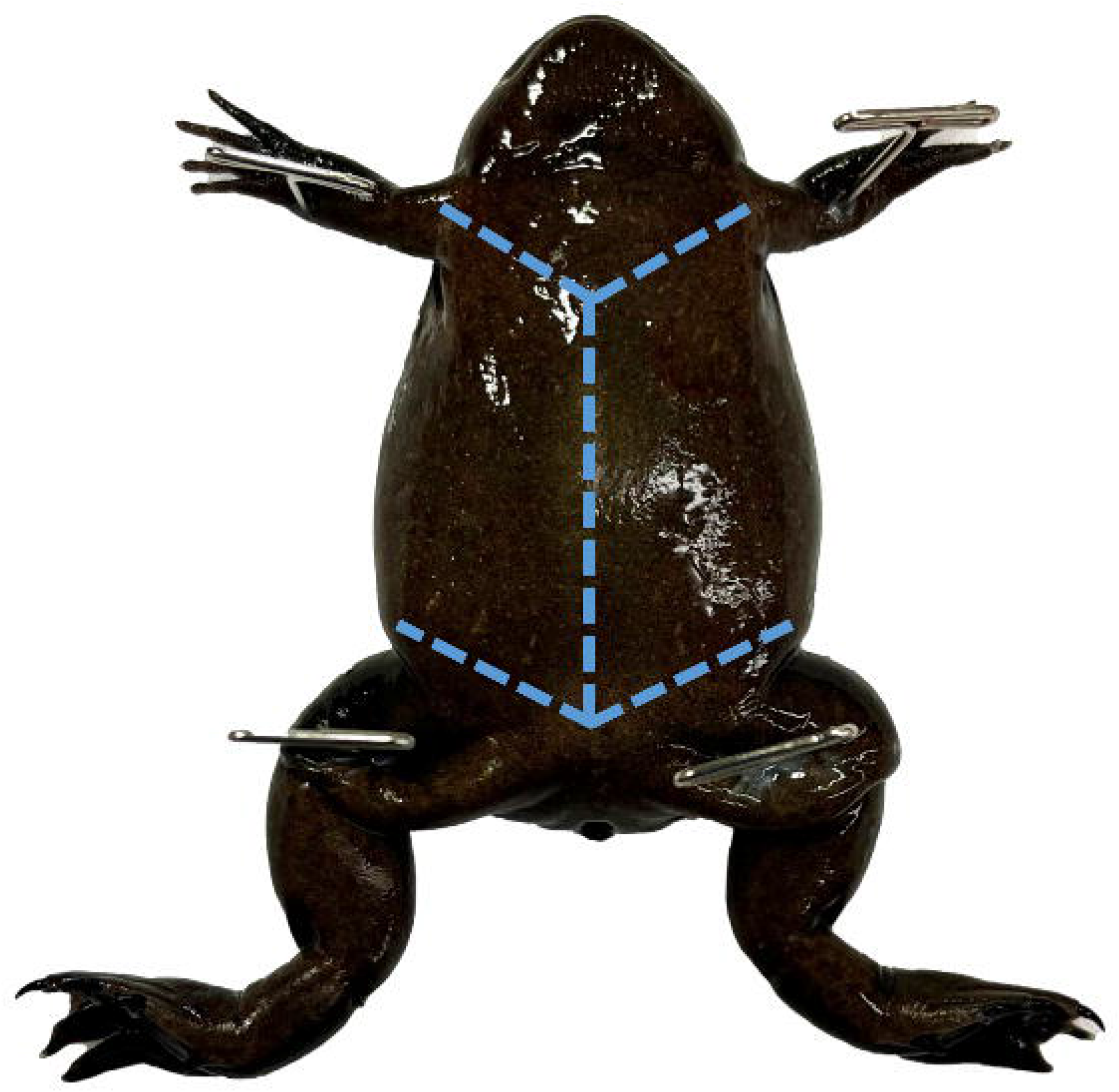
Mature female *X. tropicalis* pinned through each limb. Use toothed dissecting forceps to pull the skin near the cloaca taught to perforate it with dissection scissors and create 2 flaps as shown.

**Figure 3.**
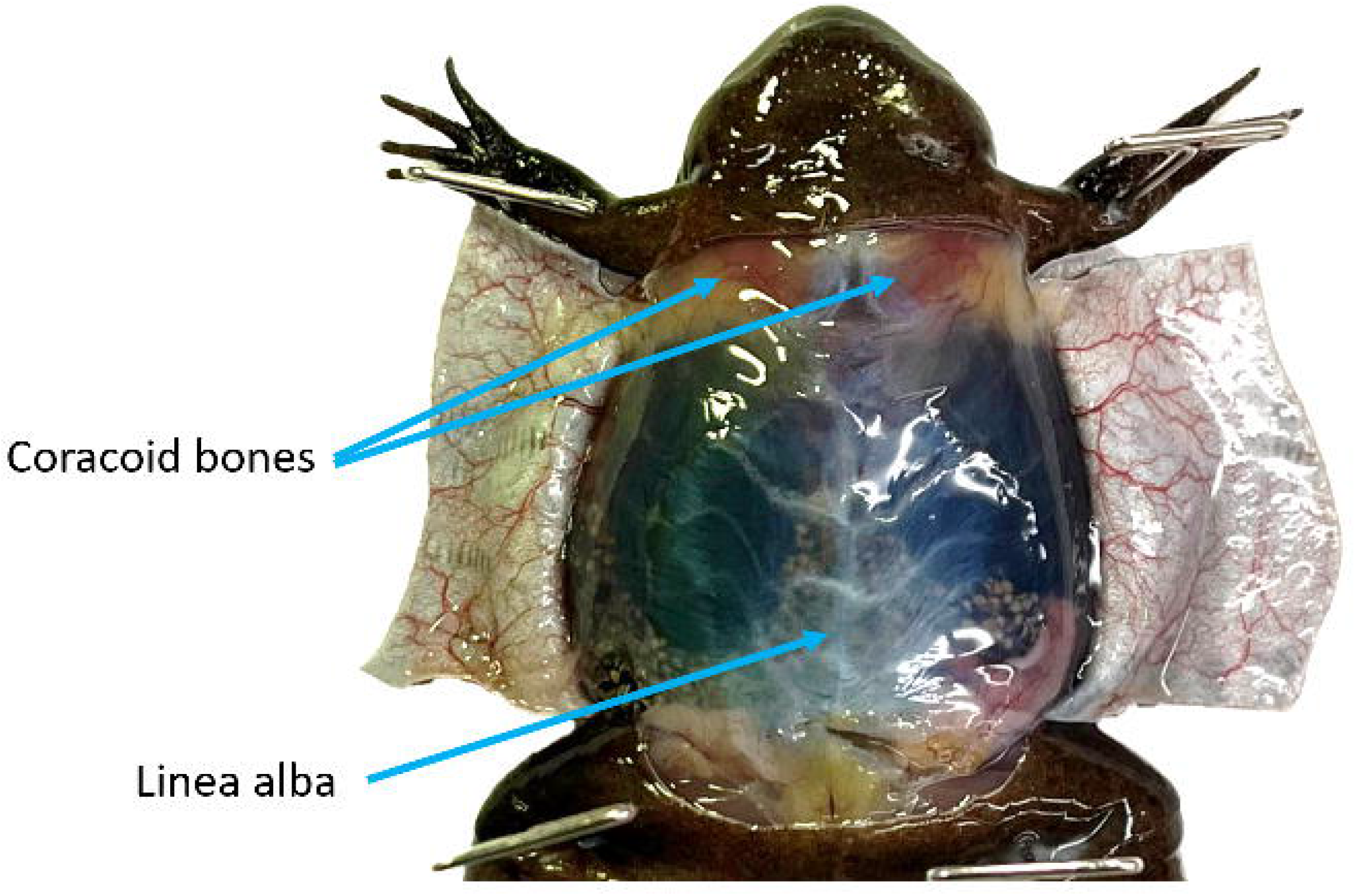
With the ventral skin open but the peritoneum intact the *linea alba* is visible. To reduce the likelihood of damaging the underlying tissues grasp the *linea alba* and pull it taught prior to cutting. The coracoid bones are visible through the peritoneum. Once the coelomic cavity has been opened these bones should be reduced to give better access to the heart.

### Perfusion

The step-by-step procedure for perfusion is defined below where we refer to respective figures and items in a separately compiled table of “troubleshooting issues”.

1. Identify the stomach and gently shift it so that it is on top of the left lobe of the liver with its vasculature visible for the duration of the procedure. Identify a lung, and grasp it by its tip. Pull it outside of the coelomic cavity and pin it through the tip, reducing unnecessary damage. Do this gently as broken blood vessels will not perfuse well (Fig. 4). Note if blood is visible within the lobe as this will affect your ability to determine the completion of the procedure.
2. If your institution allows it, take an image of the coelomic cavity to better assess perfusion efficiency and potentially identify abnormal tissues at a later date.
3. Identify the thin pericardium and pull it taught with tissue forceps. (Fig. 5) Using the tip of the iridectomy or iris scissors gently perforate it, being careful not to cut the underlying tissues. Peel the pericardium up away from the 3 chambers of the heart.
4. Use forceps to gently grasp the ventricle by its apex leaving them open enough to pass a needle through the closure. (Fig. 6)
5. Insert the needle through your forceps into the chamber of the ventricle, being careful not to perforate through it. (Fig. 7) Clamp the tissue forceps in place using a needle holder or any other clamping forceps. Clamping the needle and ventricle directly is not recommended. (Troubleshooting steps 1, 2)
6. Start the flow of the pump at approximately 5mL/min. The 3 chambers of the heart and truncus arteriosus will engorge (Fig. 8, Troubleshooting step 3).
7. Lance the right auricle (on your left), blood will pour out. You may now continue with a flow rate of 5mL/L or increase it to 10mL/min.
8. Continue until the vasculature of the stomach blanches (Troubleshooting step 5), then lance the left (your right) auricle of the heart. If the flow rate is still 5mL/min, increase the flow to approximately 10mL/min, at the time.
9. Use a transfer pipette to rinse the coelomic cavity in perfusion media to maintain visibility and better access the blood saturation of the fluid flowing from the auricles.
10. Keep the needle in place until the solution flowing from the auricles is clear (Troubleshooting step 6) and the lung has lost its red tint (Troubleshooting step 7, Fig. 9).

**Figure 4.**
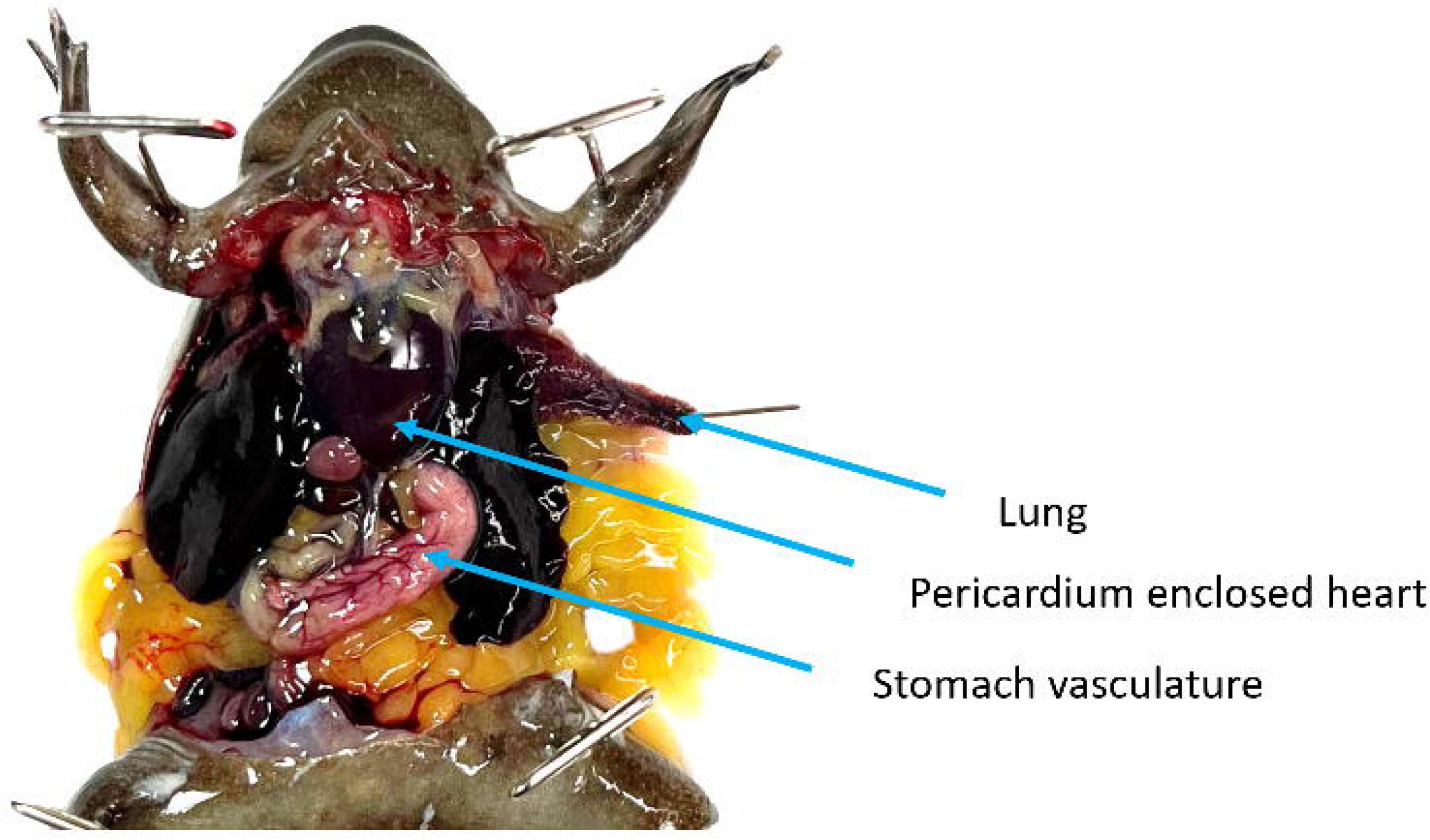
The coelomic cavity of a mature *X. tropicalis* male. The coracoid bones have been reduced providing access to the pericardium-enclosed heart. The stomach has been shifted in front of the left lobe of the liver and its vasculature is clearly visible. The left lung has been pulled out of the coelomic cavity by its tip and pinned to ensure that it does not retract during the rinsing process.

**Figure 5.**
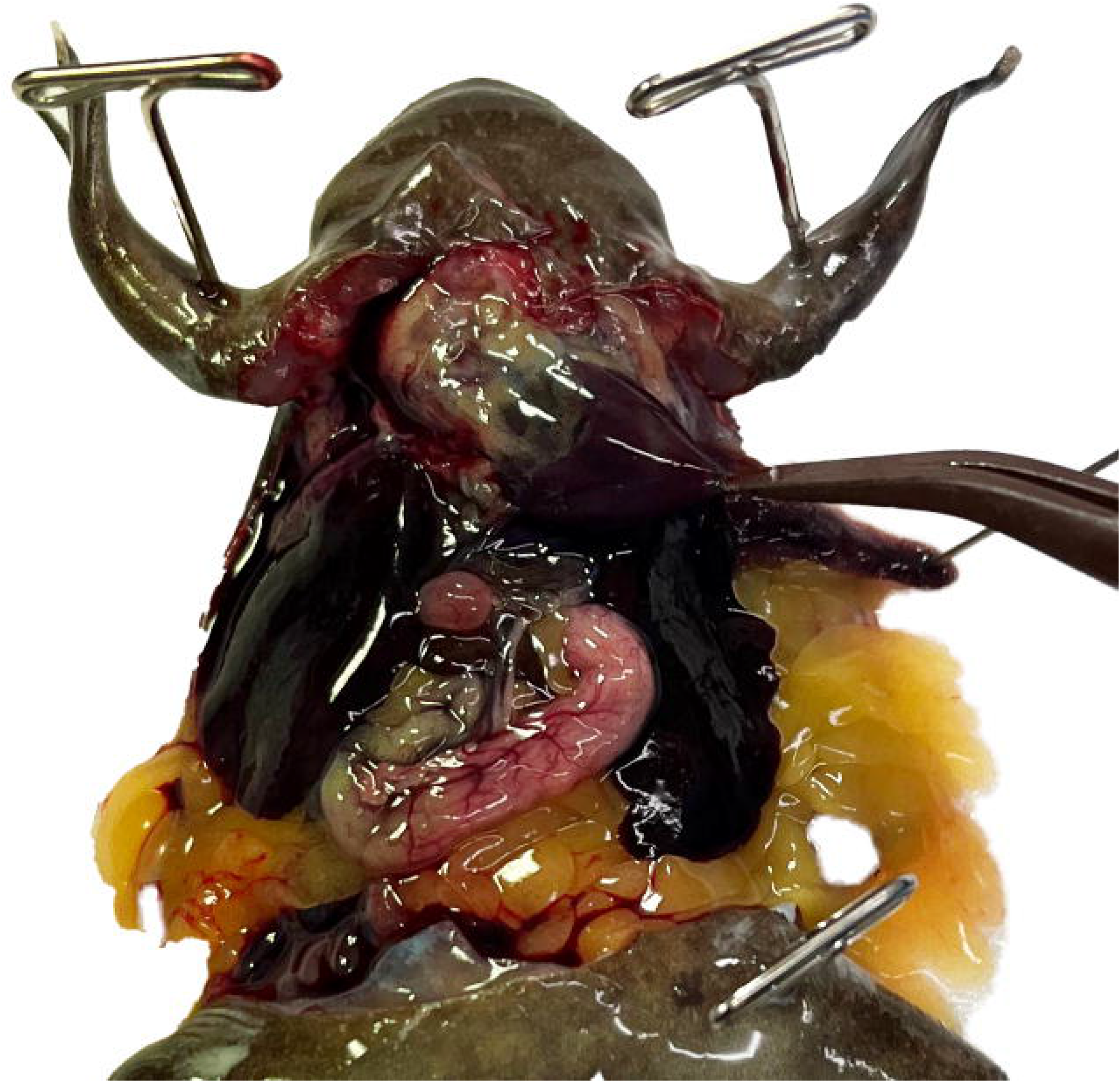
The pericardium is a thin tough membrane enclosing the heart. Using tissue forceps gently grasp the pericardium and then use the tip of iridectomy or iris scissors to perforate it. Once perforated, peel it up, away from the heart.

**Figure 6.**
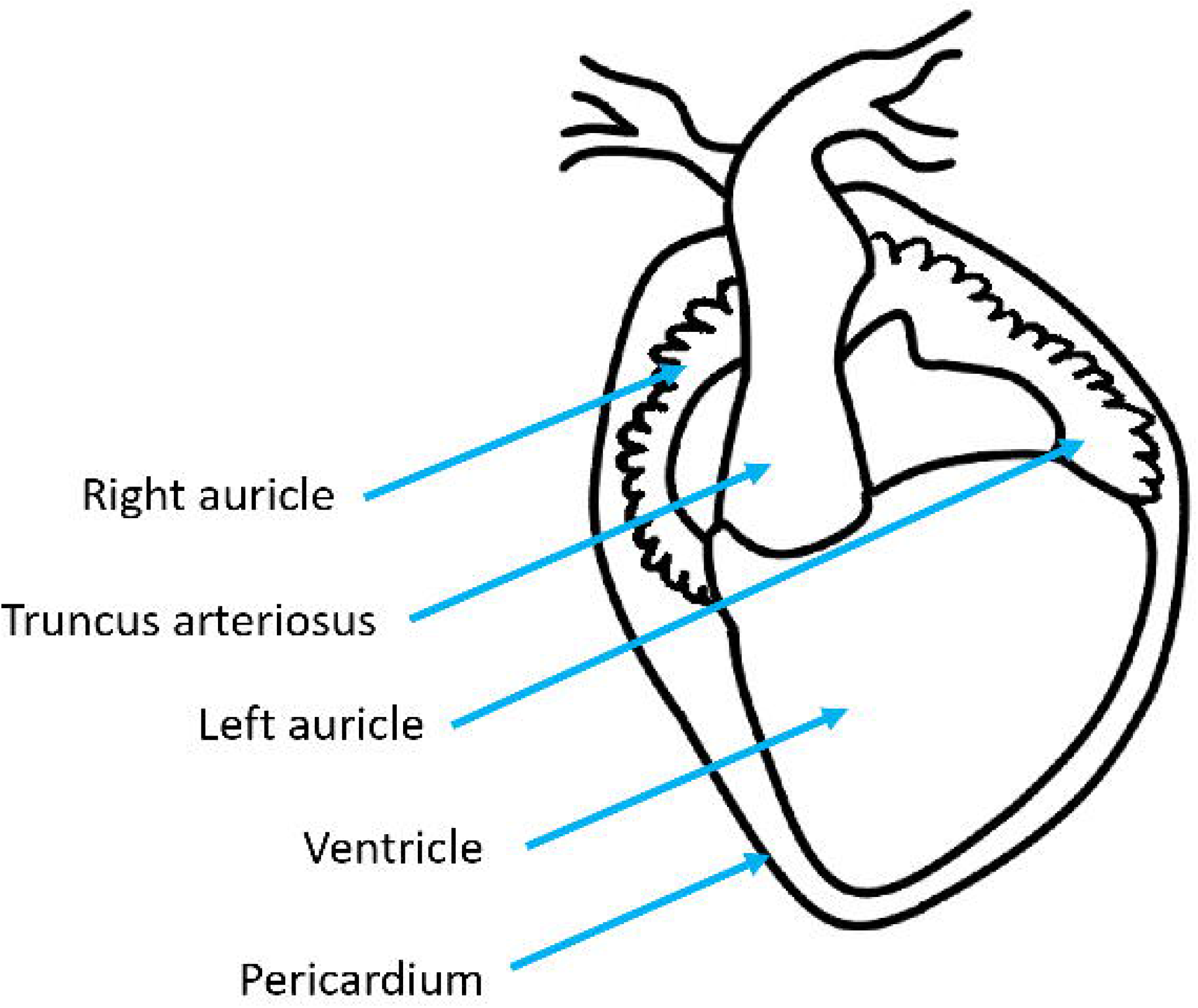
**(left)** Ventral diagram of *X. laevis* heart. **(right)** Heart diagram, with pericardium removed, showing the correct needle and clamp placement.

**Figure 7.**
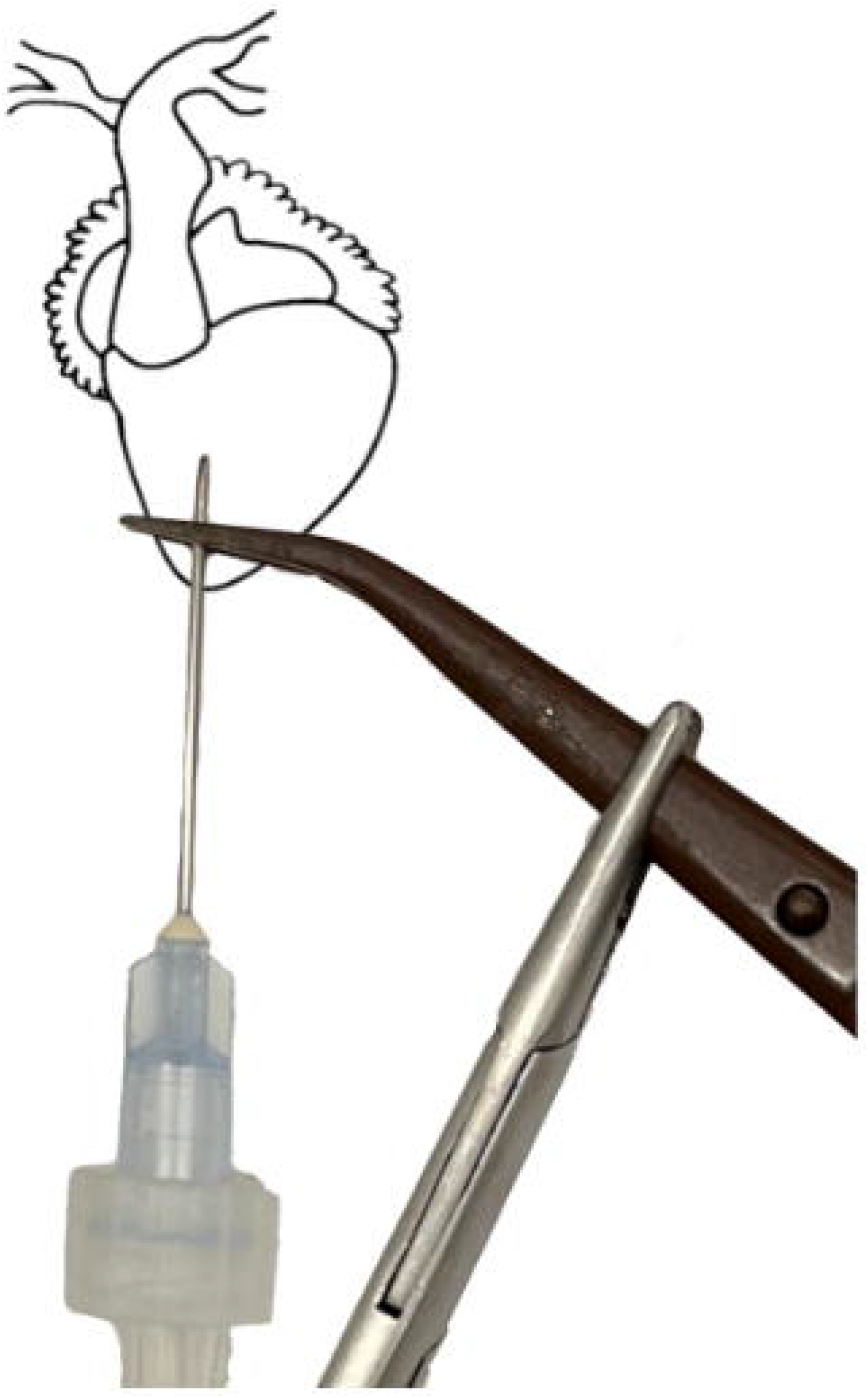
With the pericardium removed the 3 chambers of the heart and truncus arteriosus are easily visible. Use forceps to gently grasp the ventricle by its apex and then insert the needle through the forceps. Be careful not to cause unnecessary damage to the ventricle or other chambers as this will compromise the perfusion efficiency.

**Figure 8.**
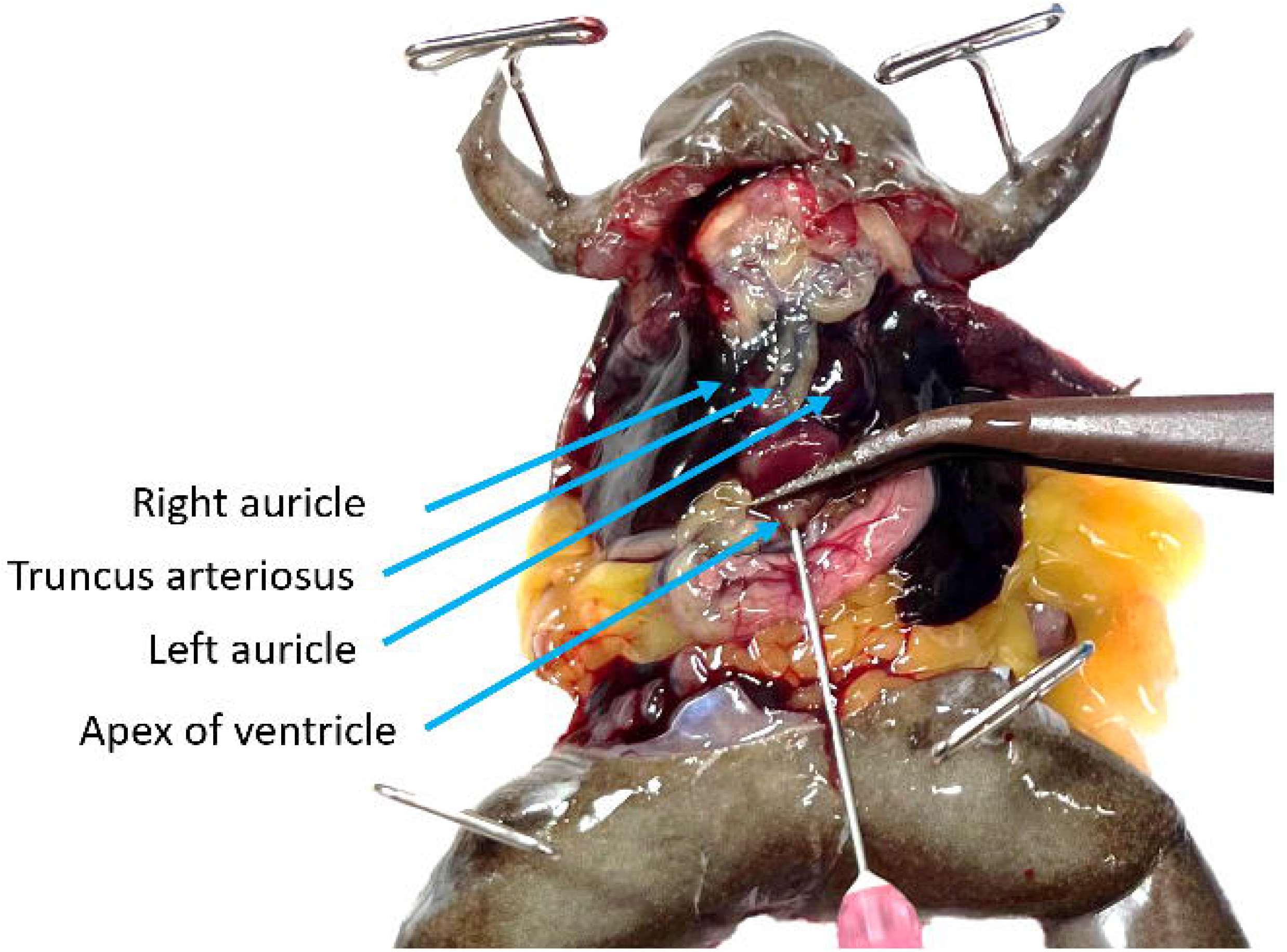
Perfusion is underway at high pressure, the right auricle has been lanced, and the ventricle, truncus arteriosus, and left auricle are visibly engorged. The stomach is blanching but both the media running from the animal and the lung tissue are heavily saturated with blood.

**Figure 9.**
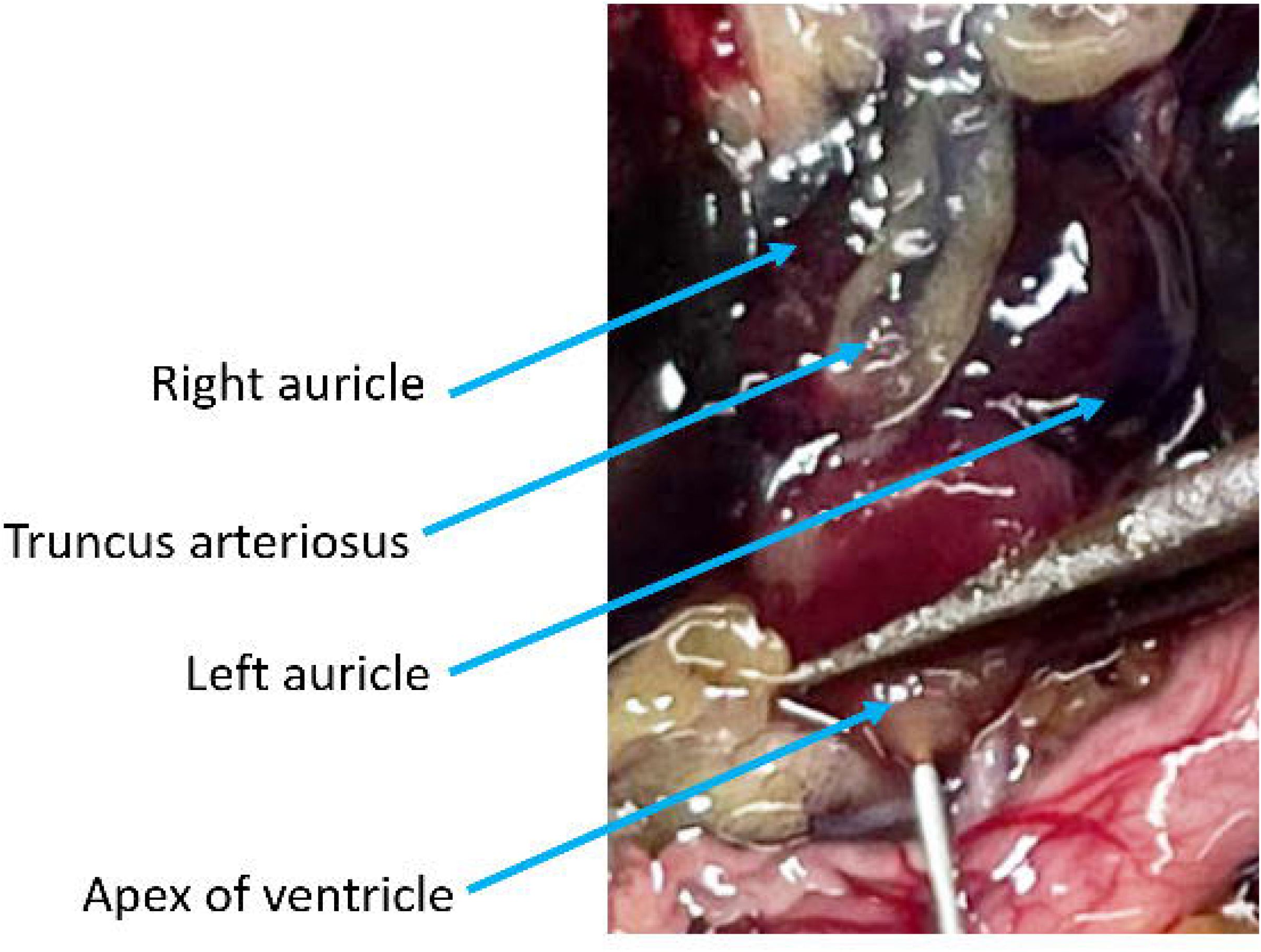
The coelomic cavity following successful rapid perfusion and rinsing. The vasculature of the stomach and other organs is no longer visible. Unless the *Xenopus* is albino the liver will remain heavily pigmented. Note that the apex of the lung still has a red tint due to blood vessels being broken from handling and pinning.

### Troubleshooting

|  | Problem | Possible cause | Solution |
| --- | --- | --- | --- |
| 1 | Needle perforates through the ventricle | Needle not adequately blunted or the heart was shifted accidentally | If possible, reclamp the needle above the perforation to isolate it (Fig. 12). If this is not possible and the perforation is small, increase the flow rate and anticipate the use of additional media.<br>If the perforation is large or cannot be isolated with the needle placed in the ventricle, perfuse into the truncus arteriosus using a blunted 27 gauge needle. Attempting a modified version of a mouse technique[2] is recommended (Fig. 12 right). Should that fail, use an axolotl technique [4] with the addition of a clamp around the truncus arteriosus and needle. |
| 2 | Needle falls out | Needle not secured prior to starting flow | Replace the needle at a greater depth and reclamp the ventricle. |
| 3 | Auricles do not swell when the pump is turned on | Pump or needle placement issues | Check to make sure the pump rotor is turning and service your pump, as needed.<br>Check that the needle is in the ventricle chamber (Troubleshooting step 1). |
| 4 | Truncus arteriosus ruptures | Air emboli or exceedingly high pressure (>20mL/minute in *X. tropicalis*, >30mL/minute in *X. laevis*) | If the truncus arteriosus ruptures you will not be able to further perfuse the animal, note incomplete perfusion. |
| 5 | Stomach doesn't blanch | vasculature may have ruptured due to air emboli or been lanced accidentally | Increase the flow up to 15mL/minute, monitor the swelling of the truncus arteriosus, if the flow rate is excessively high it will begin to bulge and may rupture.<br>Use the visibility of blood vessels in the fat bodies or other tissues to determine perfusion. |
| 6 | Tissues blanch but auricles do not run clear | Residual blood clots | Rinse any clotted blood out of the coelomic cavity using media. |
| 7 | Lung does not blanch | Broken blood vessels prior to perfusion (from handling or aspiration) | Check the lung that has not been pinned if it also has not blanched, use the color of the other tissues (Fig. 10) and the media leaving the auricles to determine complete perfusion. |
| 8 | The buffer turns red when rinsing samples other than liver tissue | Perfusion was not adequate | Give these tissues multiple rinses (until the buffer is clear) and note this, with the samples. |

## Representative results

Following successful perfusion all tissues (excluding the liver in pigmented *Xenopus*) will be distinctly lighter and less saturated with blood. Major blood vessels will become less noticeable and tissues (excluding the liver) will rinse cleanly in a buffer after being sampled. While ultimately, the successful execution of the protocol can only be confirmed by the usability of samples for the intended research project, we describe several typical issues and provide troubleshooting solutions below.

## Discussion and Remarks

This protocol describes traditional dissection techniques for accessing the coelomic cavity. Other techniques are also acceptable provided they cause minimal damage to the tissues, the heart is accessible and the lung and stomach are visible. Similarly, most of the dissection tools listed can be easily substituted with comparable items.

While we have attempted to explore the efficacy of this procedure to the extent possible within the limits of a single research lab, the results may vary depending on particular environments and research circumstances. One interesting aspect of blood perfusion procedure that remained outside of the scope of this paper is how this procedure compares to alternative ways of perfusion for animals that underwent surgery. Another unexplored variable is how blood perfusion would work in very young animals or animals of advanced age where vasculature might be excessively fragile. Below, we provide several additional remarks we hope will facilitate the application of our protocol.

If perfusion efficiency takes precedence over rapid perfusion, adapting an axolotl technique is recommended [4]. Note that the Saltman et al. paper uses the term aorta to refer to the truncus arteriosus.

Note that the duration of the procedure and volume of media used is dependent on a number of variables. Generally, *X. tropicalis* males take between 2 and 3 minutes to successfully perfuse with 15-25 mL of media while *X. tropicalis* females take between 3 and 4 minutes with 25-40mL of media. Significantly more animal-to-animal variation was found when perfusing *X. laevis*.

Naturally, it is much easier to assess perfusion efficiency in albino animals. The difference is especially apparent in the lung and liver tissues (Fig. 10). Thus the use of albinos is recommended, especially when first attempting perfusion or undergoing training.

**Figure 10.**
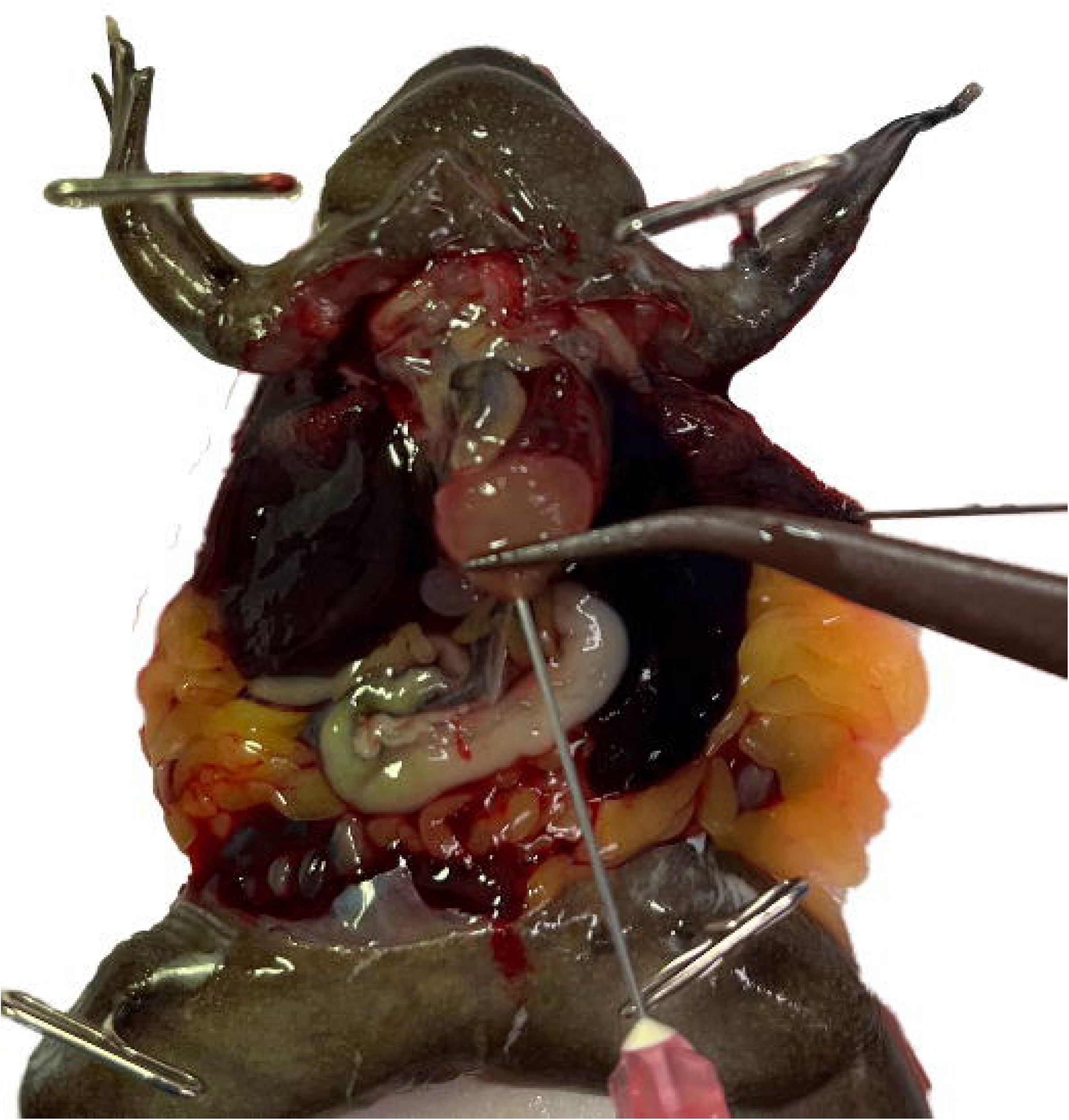
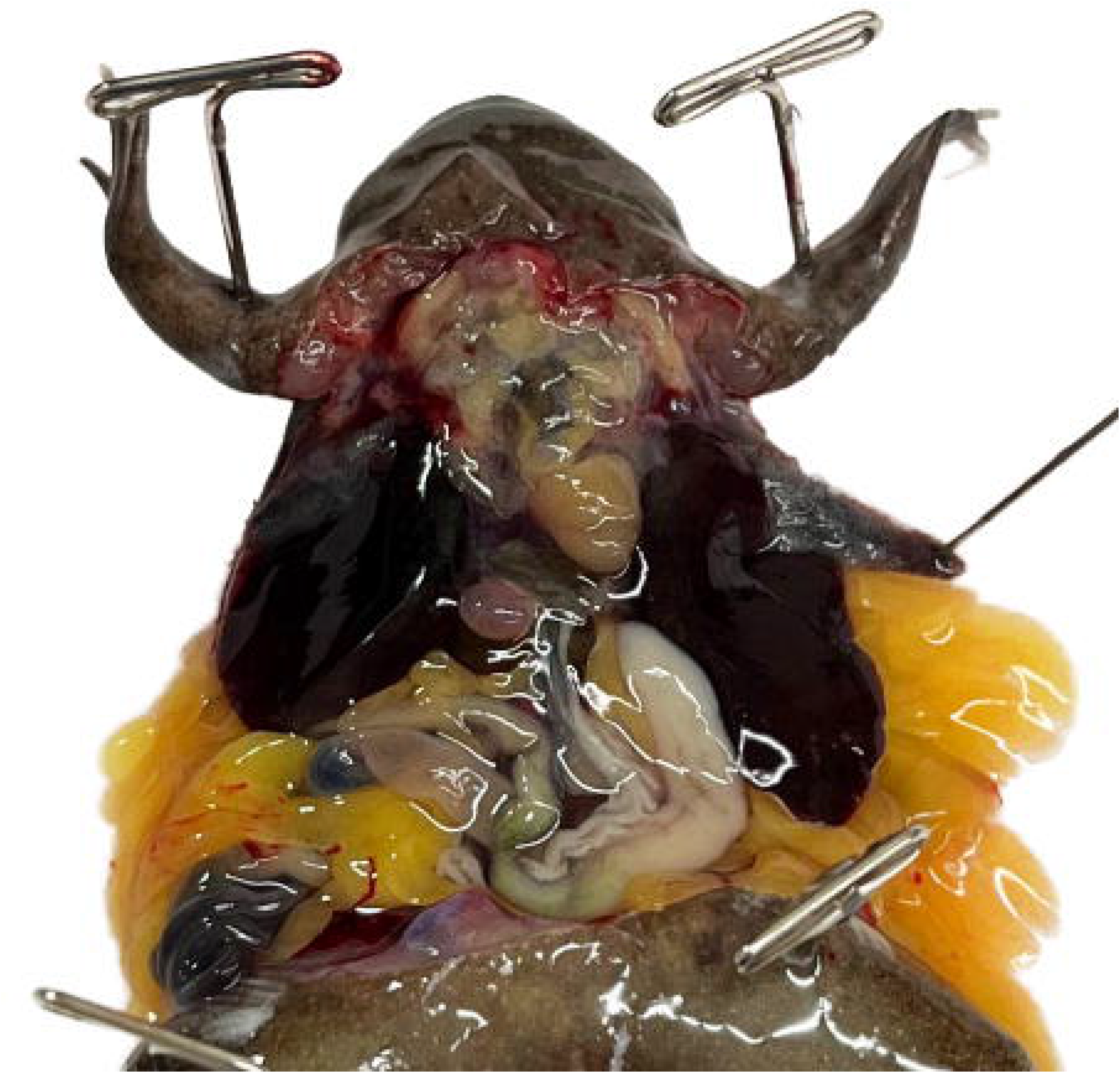
*X. laevis* fat body lobe sampled before **(left)** perfusion and a fat body lobe of the same individual, sampled after **(right)** perfusion.

**Figure 11.**
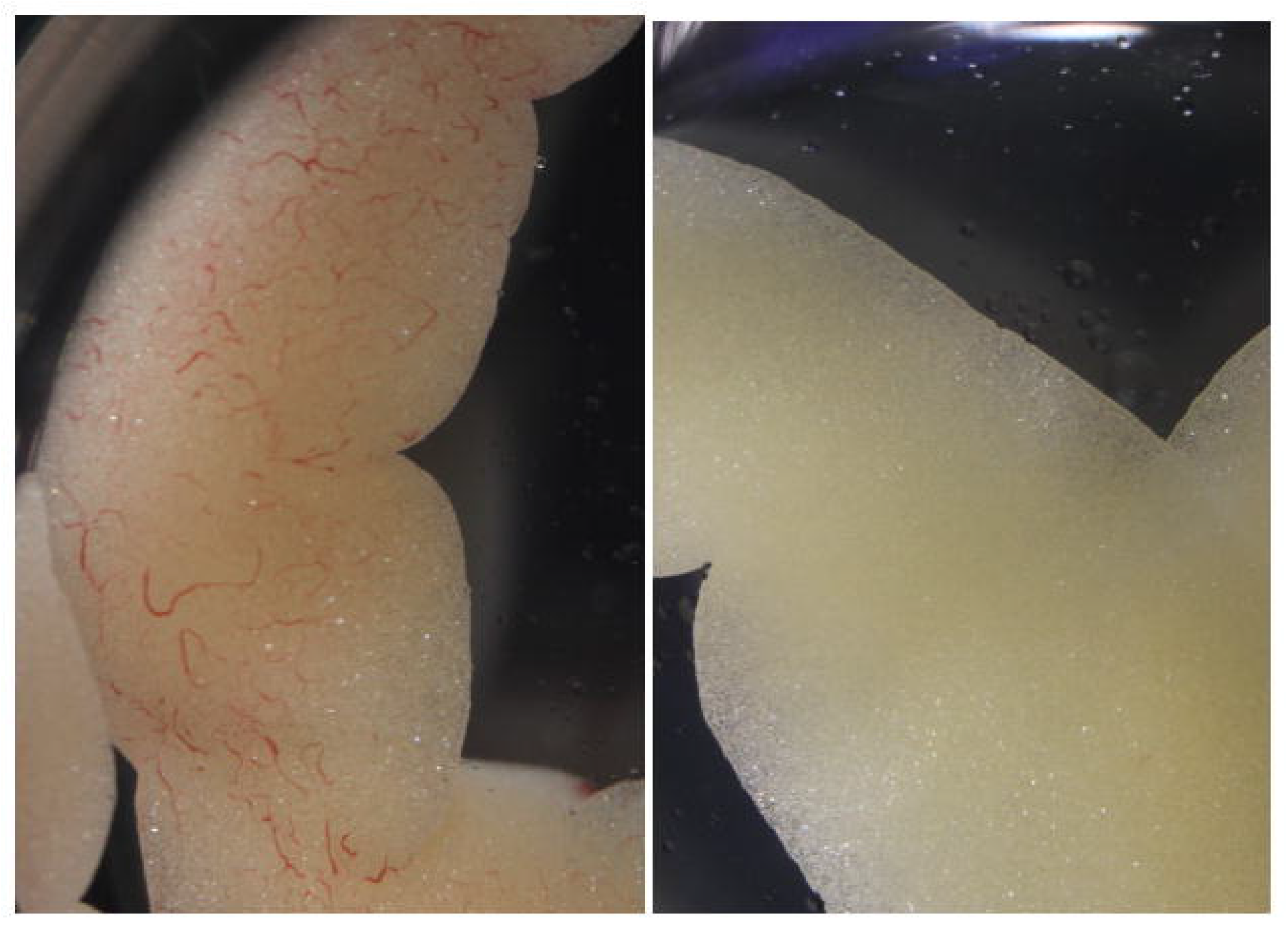
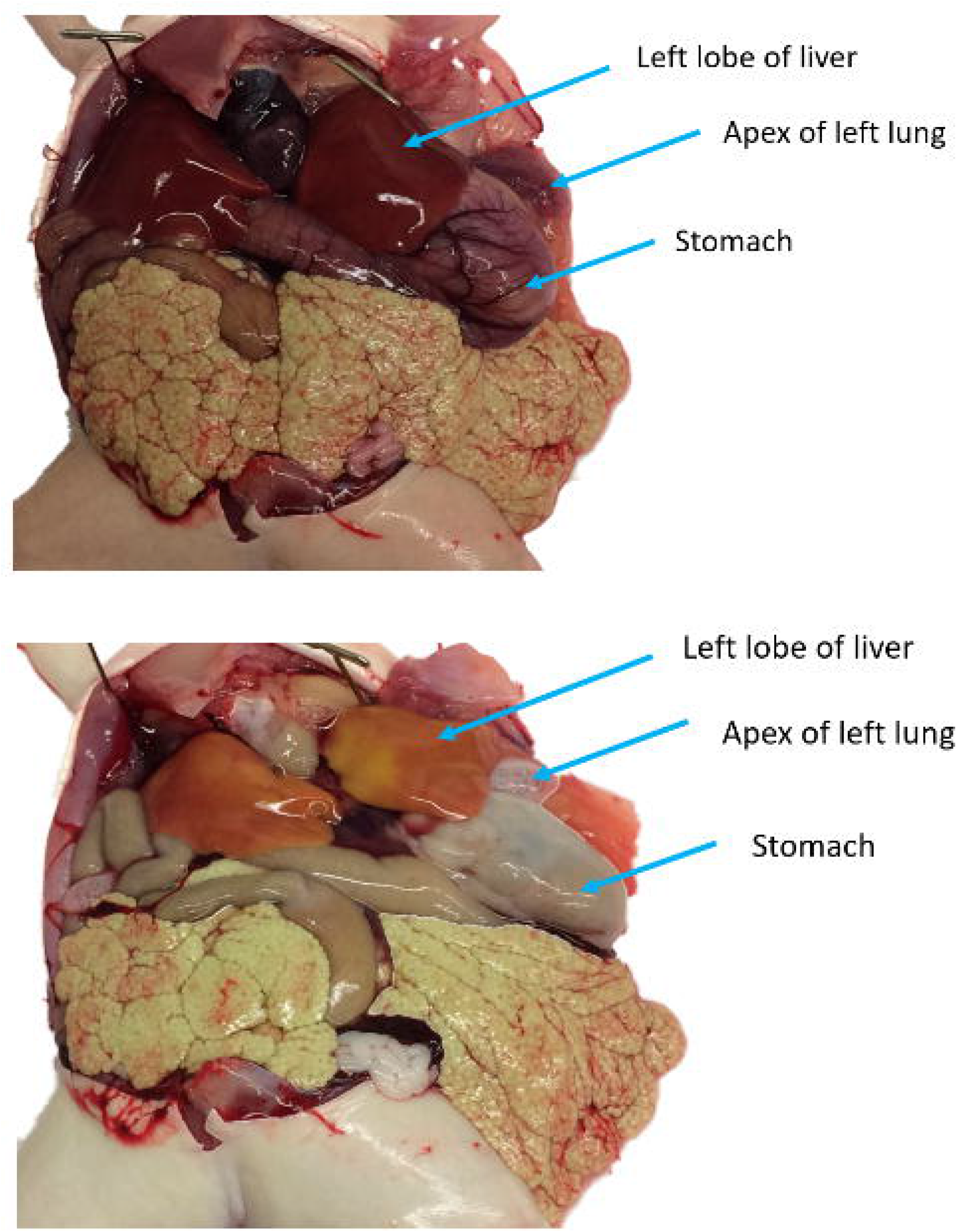
An albino *X. laevis* both before **(top)** and after **(bottom)** rapid perfusion. The albinism makes it easier to determine the proficiency of the perfusion than it would be in a pigmented animal. This is especially apparent in the lung and liver tissues.

**Figure 12.**
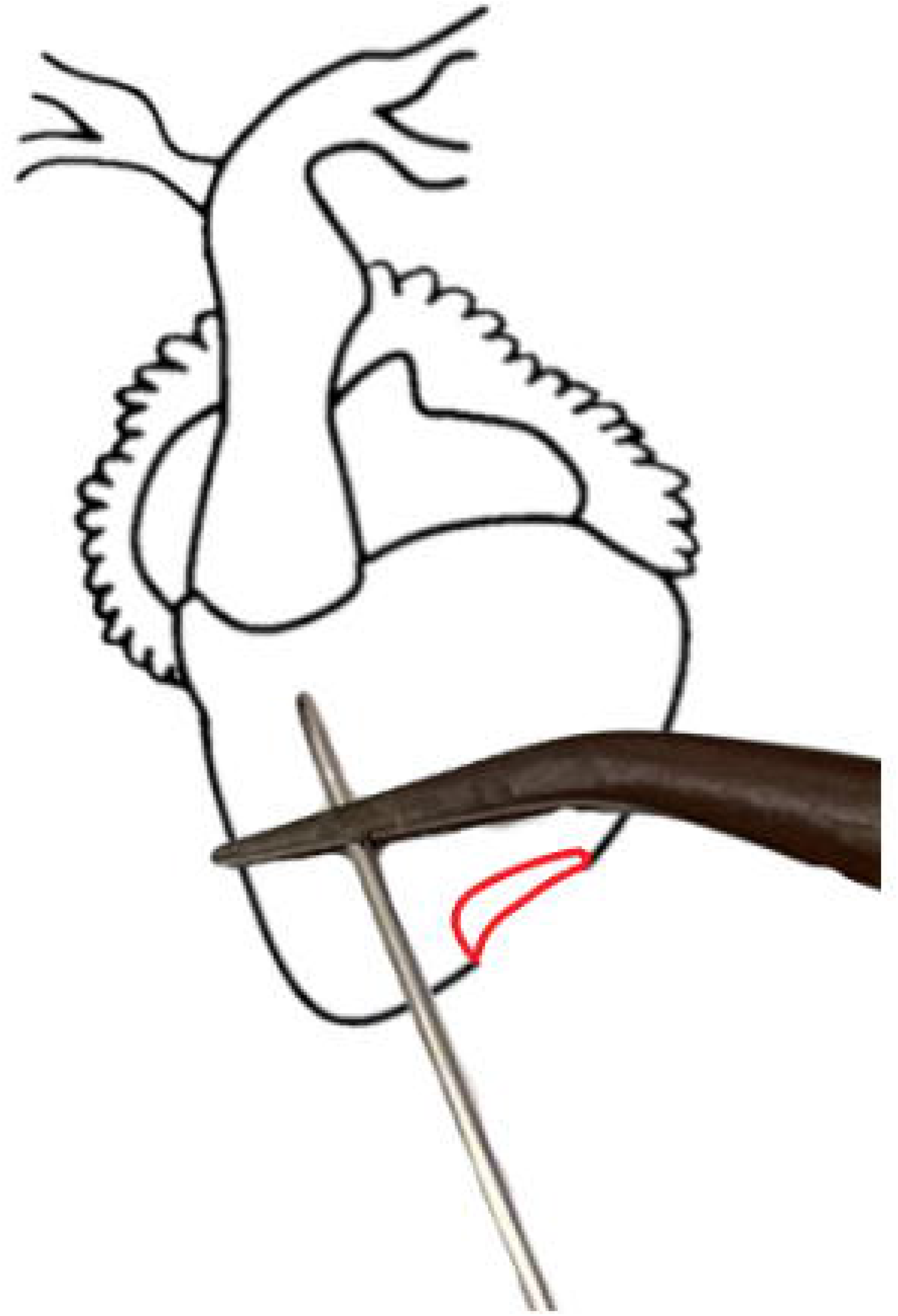
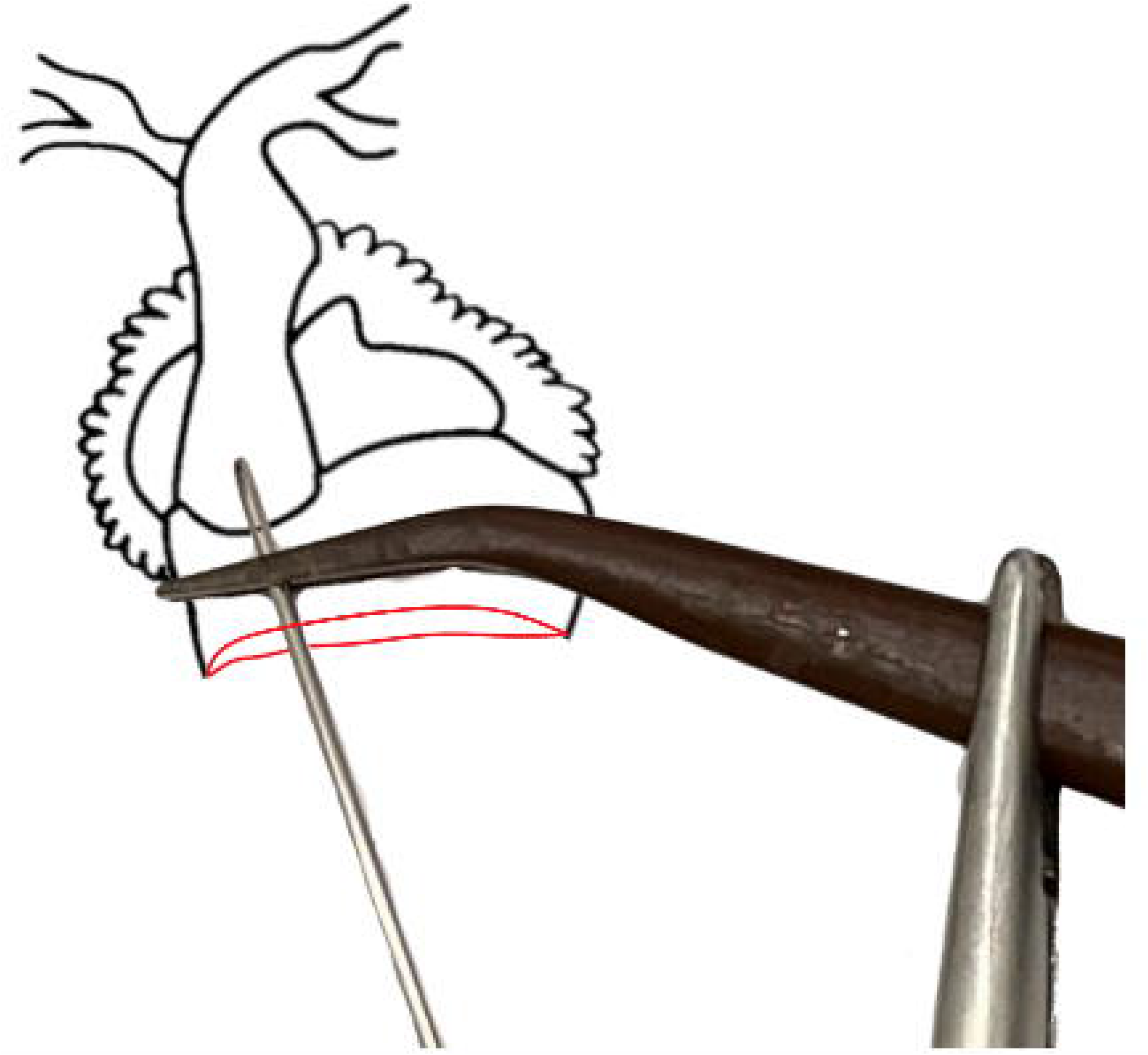
Troubleshooting diagram of *X. laevis* heart. **(left)** The ventricle has perforation (in red) this perforation is isolated by the forceps and will not affect perfusion efficiency. **(right)** A heart with a severely damaged ventricle. The needle can be guided into the truncus arteriosus and clamped in place. It is especially important to ensure that the needle is well-blunted when using this technique [2].

By adjusting the flow rate and needle size the protocol is adaptable for all species of *Xenopus*. Owing to the homology in heart anatomy and blood circulation between *Xenopus* and most other amphibians [13] as well as non-crocodilian reptiles this technique may be modified for the full-body rapid perfusion of other models with 3-chambered hearts [14]. Were there a non-crocodilian reptile model that exclusively required the perfusion of one of the two aortic arches, other protocols are recommended [15].

## Disclosures

The authors declare no competing interests.

## Acknowledgments

This work was supported by NIH’s OD R24 grant ODO31956 and NICHD R01 grant HD073104. We thank Darcy Kelly for helpful discussions and initial input on this protocol as well as Samantha Jalbert and Jill Ralston for their assistance and support.

---

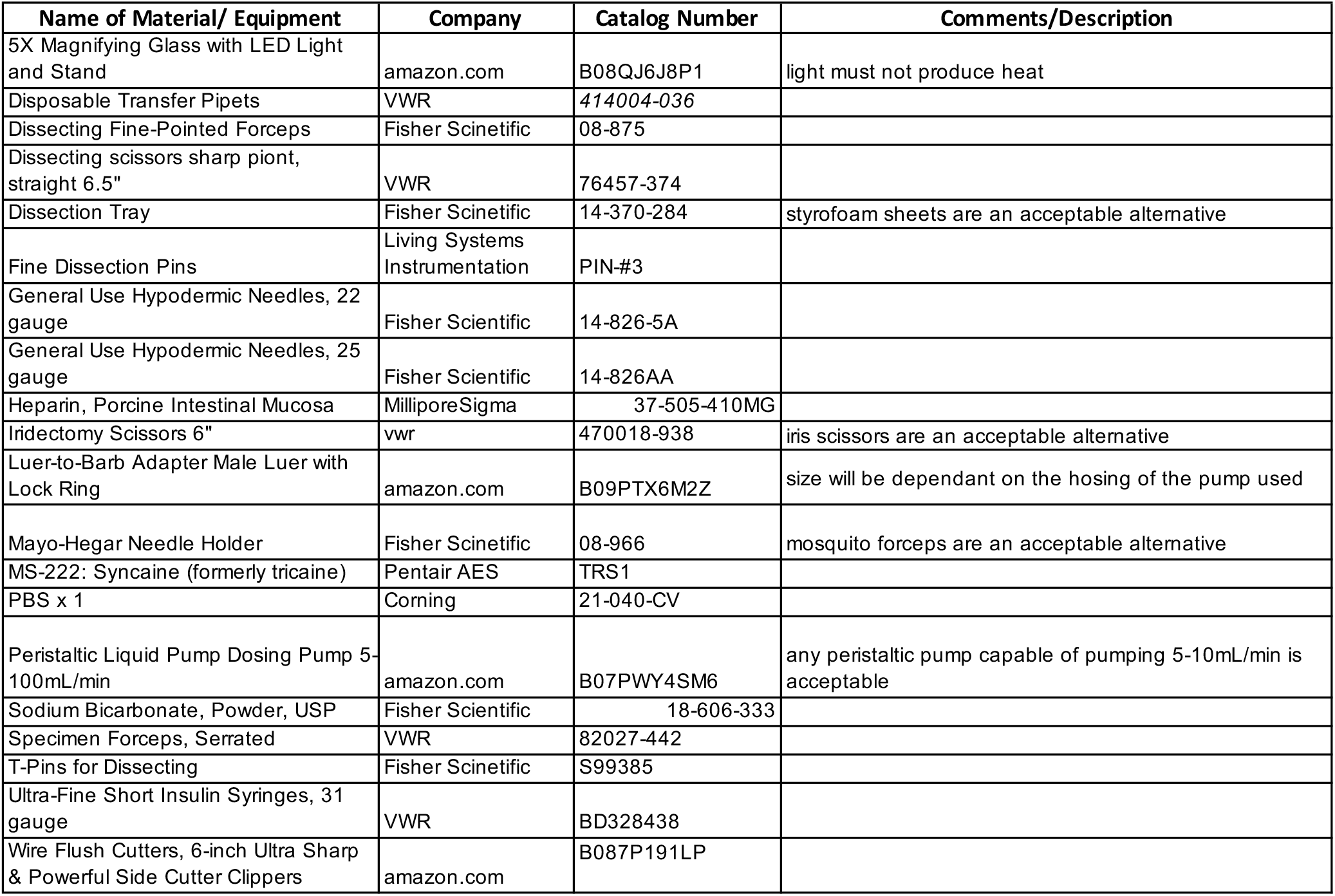

## Notes

### Competing Interest Statement

The authors have declared no competing interest.

